# A computational model tracks whole-lung *Mycobacterium tuberculosis* infection and predicts factors that inhibit dissemination

**DOI:** 10.1101/713701

**Authors:** Timothy Wessler, Louis R. Joslyn, H. Jacob Borish, Hannah P. Gideon, JoAnne L. Flynn, Denise E. Kirschner, Jennifer J. Linderman

## Abstract

*Mycobacterium tuberculosis* (Mtb), the causative infectious agent of tuberculosis (TB), kills more individuals per year than any other infectious agent. Granulomas, the hallmark of Mtb infection, are complex structures that form in lungs, composed of immune cells surrounding bacteria, infected cells, and a caseous necrotic core. While granulomas serve to physically contain and immunologically restrain bacteria growth, some granulomas are unable to control Mtb growth, leading to bacteria and infected cells leaving the granuloma and disseminating, either resulting in additional granuloma formation (local or non-local) or spread to airways or lymph nodes. Dissemination is associated with development of active TB. It is challenging to experimentally address specific mechanisms driving dissemination from TB lung granulomas. Herein, we develop a novel hybrid multi-scale computational model, *MultiGran,* that tracks Mtb infection within multiple granulomas in an entire lung. *MultiGran* follows cells, cytokines, and bacterial populations within each lung granuloma throughout the course of infection and is calibrated to multiple non-human primate (NHP) cellular, granuloma, and whole-lung datasets. We show that *MultiGran* can recapitulate patterns of *in vivo* local and non-local dissemination, predict likelihood of dissemination, and predict a crucial role for multifunctional CD8+ T cells and macrophage dynamics for preventing dissemination.

**Author Summary:** Tuberculosis (TB) is caused by infection with *Mycobacterium tuberculosis* (Mtb) and kills 3 people per minute worldwide. Granulomas, spherical structures composed of immune cells surrounding bacteria, are the hallmark of Mtb infection and sometimes fail to contain the bacteria and disseminate, leading to further granuloma growth within the lung environment. To date, the mechanisms that determine granuloma dissemination events have not been characterized. We present a computational multi-scale model of granuloma formation and dissemination within primate lungs. Our computational model is calibrated to multiple experimental datasets across the cellular, granuloma, and whole-lung scales of non-human primates. We match to both individual granuloma and granuloma-population datasets, predict likelihood of dissemination events, and predict a critical role for multifunctional CD8+ T cells and macrophage-bacteria interactions to prevent infection dissemination.

## Introduction

Tuberculosis (TB) kills more individuals per year than any other infectious disease and treatment remains a global challenge (1). Only a small fraction (5-10%) of those infected with *Mycobacterium tuberculosis* (Mtb) develop active symptomatic disease (2), while the remainder control but do not eliminate the infection, which is termed latent TB (LTBI). A hallmark of Mtb infection is the presence of lung granulomas (lesions), collections of immune cells that surround Mtb in an effort to contain and control an infection. Multiple granulomas can be present in humans and non-human primates (NHPs). In NHPs, each granuloma is initiated by a single bacillus (3). Of key importance is that each granuloma within an individual has its own independent trajectory behavior. For example, the immune response in some granulomas eliminates all bacteria, resulting in sterilization. In other granulomas, immune cells only contain Mtb growth, resulting in stable granulomas that may persist for decades (4). If Mtb growth is not contained, however, granulomas can grow and/or spread, allowing for dissemination of bacteria across the lungs leading to the formation of new granulomas, spread to the airways resulting in transmission of infection through aerosolized bacteria, and possibly death of the host if not treated. Understanding the collective behavior of granulomas within lungs leading to dissemination events is critical to the ultimate goal of controlling the global TB epidemic.

It is difficult to experimentally address specific mechanisms operating within lungs that drive different granuloma outcomes in primates, although it is known through interventional studies that certain factors, such as TNF, CD4+ T cells, and CD8+ T cells are important in controlling early and established Mtb infection (5–8). As a complementary approach, mathematical modeling can generate hypotheses that can then be tested experimentally. Several mathematical and computational models for Mtb infection have been developed to explore the contributions of the innate and adaptive immune responses to granuloma formation and function (9–20). These models are informed by studies in humans and in animal models of infection, especially NHPs, rabbits, pigs, and mice (21). In particular, *GranSim,* our computational model that allows simulation of the formation and function of a single granuloma using a hybrid agent-based model framework, has offered strategies for drug treatment and vaccine development (12,14,22–24). *GranSim*, which considers thousands of cells and bacteria as “agents” in the simulation and tracks diffusion of multiple immune mediators (e.g., cytokines), is computationally intensive, limiting our ability to simultaneously simulate multiple granulomas present in an entire lung during infection. In contrast, Prats et al. (18) utilized a bubble model to demonstrate the importance of local inflammation, dissemination, and coalescence of lesions as key factors leading to active TB, but did not specifically model events at the granuloma scale. However, following the formation of individual granulomas, the dissemination of those granulomas across the lungs over time, and, importantly, tracking events at the granuloma scale could provide an important window into infection dynamics and could lead to new insights for prevention or treatment.

In order to study the formation of new granulomas after initial establishment of infection, referred to as dissemination, the evolution of individual granulomas must be captured over time. Recently, research on Mtb-infected NHPs provided data on disseminating granulomas (25). Of all animal models used to study Mtb infection, NHPs are most relevant to human TB disease because they recapitulate the full spectrum of clinical outcomes and pathologies seen in humans (26). From PET CT imaging, the emergence of new granulomas was tracked and recorded. The authors genetically matched Mtb barcodes, assigned each inoculation Mtb a unique barcode ID, and associated each granuloma identified in the temporal PET CT images with the Mtb barcodes inside that granuloma (Figure 1). By identifying Mtb barcodes that were present in multiple granulomas, they were able to distinguish disseminating from non-disseminating granulomas. When identifying multiple bacterial barcodes within a single granuloma, it is surmised a merger of granulomas took place. While Martin et al. showed these distinctions, the mechanisms that lead to granuloma clustering or dissemination remain unanswered. We address these open questions using a hybrid computational-mathematical modeling framework.

**Fig 1.**
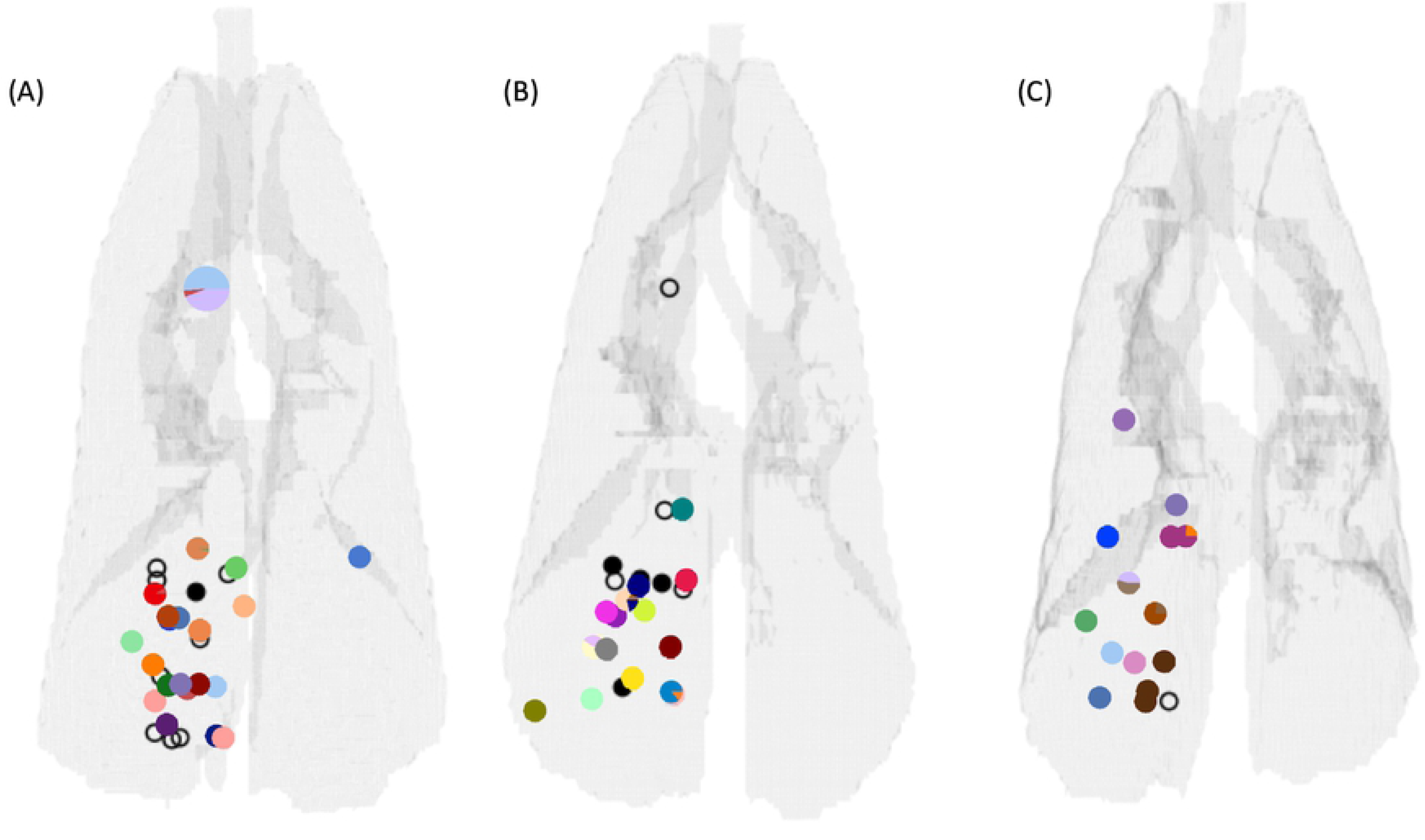
Three NHP lung maps illustrating the positioning of pulmonary granulomas and thoracic lymph nodes (data previously published in Martin et al. (22)). Gray outlines denote the extent of the lungs, bronchial tubes, and trachea. Small markers superimposed on the outlines represent the positions of pulmonary granulomas, while larger markers denote lymph nodes. Colors denote unique barcode tags. Some samples had more than one barcode tag present, and often these were doublet granulomas (i.e., two granulomas too close in proximity to distinguish at necropsy) and so are marked with a pie chart showing the relative abundance of each barcode tag. The black markers represent pulmonary granulomas for which no barcode tags were found. Filled black markers are granulomas which grew bacteria upon plating but barcodes could not be determined for technical reasons, while open markers are granulomas that did not grow bacteria upon plating (sterile).

Herein, we develop a novel multi-scale hybrid model, *MultiGran,* to track Mtb infection at the scale of the entire lung, including capturing multiple granulomas and their individual outcomes as well as the formation of new granulomas. *MultiGran* is an agent-based model that follows cells, cytokines, and bacterial populations across multiple lung granulomas throughout the course of infection. Each granuloma is now formulated as a single agent, and each agent contains within it a system of non-linear ordinary differential equations (ODEs) that capture individual granuloma dynamics. *MultiGran* follows the steps observed through the course of Mtb infection: (1)*initial granuloma establishment* with Mtb that have been virtually barcoded and placed within the lung environment, (2) *granuloma development* across time, (3) the possibility of *granuloma dissemination* with barcoded bacteria moving to a new location, and (4) *granuloma merging* by granulomas that have formed close together and whose individual boundaries are indistinguishable, or those that grow in size and thus merge into a granuloma cluster (that may have multiple barcoded bacteria IDs). We use *MultiGran* to address three outstanding questions about dissemination: what mechanisms are consistent with granuloma dissemination and merging patterns seen *in vivo*? What is the likelihood of a granuloma to disseminate? Can we predict factors that lead to dissemination?

## Methods

### Ethics Statement

All experimental manipulations, protocols, and care of the animals were approved by the University of Pittsburgh School of Medicine Institutional Animal Care and Use Committee (IACUC). The protocol assurance number for our IACUC is A3187-01. Our specific protocol approval numbers for this project are 1280653, 12126588, 11110045, 19024273, 15066174, 16017309 and 18124275. The IACUC adheres to national guidelines established in the Animal Welfare Act (7 U.S.C. Sections 2131 - 2159) and the Guide for the Care and Use of Laboratory Animals (8^th^ Edition) as mandated by the U.S. Public Health Service Policy.

All macaques used in this study were housed at the University of Pittsburgh in rooms with autonomously controlled temperature, humidity, and lighting. Animals were singly housed in caging at least 2 square meters apart that allowed visual and tactile contact with neighboring conspecifics. The macaques were fed twice daily with biscuits formulated for NHPs, supplemented at least 4 days/week with large pieces of fresh fruits or vegetables. Animals had access to water *ad libitem*. Because our macaques were singly housed due to the infectious nature of these studies, an enhanced enrichment plan was designed and overseen by our nonhuman primate enrichment specialist. This plan has three components. First, species-specific behaviors are encouraged. All animals have access to toys and other manipulata, some of which will be filled with food treats (e.g., frozen fruit, peanut butter). These are rotated on a regular basis. Puzzle feeders foraging boards, and cardboard tubes containing small food items also are placed in the cage to stimulate foraging behaviors. Adjustable mirrors accessible to the animals stimulate interaction between animals. Second, routine interaction between humans and macaques are encouraged. These interactions occur daily and consist mainly of small food objects offered as enrichment and adhere to established safety protocols. Animal caretakers are encouraged to interact with the animals (by talking or with facial expressions) while performing tasks in the housing area. Routine procedures (e.g., feeding, cage cleaning) are done on a strict schedule to allow the animals to acclimate to a routine daily schedule. Third, all macaques are provided with a variety of visual and auditory stimulation. Housing areas contain either radios or TV/video equipment that play cartoons or other formats designed for children for at least 3 hours each day. The videos and radios are rotated between animal rooms so that the same enrichment is not played repetitively for the same group of animals.

All animals are checked at least twice daily to assess appetite, attitude, activity level, hydration status, etc. Following Mtb infection, the animals are monitored closely for evidence of disease (e.g., anorexia, weight loss, tachypnea, dyspnea, coughing). Physical exams, including weights, are performed on a regular basis. Animals are sedated prior to all veterinary procedures (e.g., blood draws) using ketamine or other approved drugs. Regular PET/CT imaging is conducted on most of our macaques following infection and has proved very useful for monitoring disease progression. Our veterinary technicians monitor animals especially closely for any signs of pain or distress. If any are noted, appropriate supportive care (e.g., dietary supplementation, rehydration) and clinical treatments (analgesics) are given. Any animal considered to have advanced disease or intractable pain or distress from any cause is sedated with ketamine and then humanely euthanatized using sodium pentobarbital.

### Experimental dataset

Experimental data specifically for this study were obtained from seven cynomolgus macaques (*Macaca fascicularis*), infected with low dose Mtb (Erdman strain, ∼10 CFU by bronchoscopic instillation) as previously described (27–29). Infection was confirmed by PET CT imaging. PET CT scans were performed monthly to quantify new granuloma formation or clustering, as well as disease progression. Necropsy was performed as previously described (28, 29). Briefly, an ^18^F-FDG PET-CT scan was performed on every animal 1-3 days prior to necropsy to measure disease progression and identify individual granulomas and other pathologies as described (27–30); this scan was used as a map for identifying individual lesions. At necropsy, each granuloma or other pathologies from lung and mediastinal lymph nodes were obtained for histological analysis, bacterial burden, and immunological studies, including flow cytometry, as previously described (27–30). For bacterial burden, each granuloma homogenate was plated onto 7H11 medium, and the CFU were enumerated 21 days later to determine the number of bacilli in each granuloma (27, 29).

To calibrate the individual granuloma computational model, we excised granulomas from macaques that were infected for 3 weeks (n=2), 5 weeks (n=2), 7 weeks (n=2) and 9 weeks (n=1). In addition, an animal without Mtb infection was also included in this study as a control. To obtain accurate cell number measurements, enzymatic digestion (Tumor dissociation kit, human; Miltenyi Biotec) was performed on excised granulomas using gentleMACS octo dissociator. The single cell suspension obtained by enzymatic digestion was processed for bacterial burden and cell numbers enumeration (27). Single cell suspensions of individual granulomas were stained with cell surface antibodies to enumerate T cells (CD3) and macrophages (CD11b). The cells were further stained intracellularly with Calprotectin antibody to exclude CD11b+Calprotectin+ cells from macrophage population. Flow cytometry and data acquisition was performed using BD LSRII and analysis was performed using Flowjo Software v10 (27).

In addition, bacterial burden data of 623 granulomas from 38 NHP that were controls in other studies (previously published (20,27,31–33) and ongoing studies) at University of Pittsburgh (Flynn Lab) were included for evaluation. The timing of infection depended on the particular study (Table of CFU values and tables of cell counts located at http://malthus.micro.med.umich.edu/labmovies/MultiGran/) Table: gran-cfu-cyno-size) and ranged from 4-17 weeks post Mtb infection.

### Non-human primate lung lattice data

To create a virtual lung that replicates an NHP lung, we used a CT scan of an uninfected NHP to model the 3-dimensional lung space. Binary images mapping the cross section of the lungs were created for each CT slice by segmentation of CT image values below −320 Houndsfield units. The individual slices were then stacked into an array, and a polygon mesh outlining the lung volume was generated using the marching_cubes_classic function in the open source Python scikit-image package (v 0.14.1, (34)).

### Identifying granuloma distributions in Non-human primate lungs

To allow us to test whether the distribution of granulomas in our virtual lungs matched that observed in NHP lungs, we refer to the distribution of granulomas arising from barcoded bacteria derived from our previously published data in Martin et al. (25). In that study, four cynomolgus macaques were infected with 11+/- 5 CFU barcoded Mtb Erdman. Barcoded libraries were generated where each bacterium has a different random 7-mer along with one of three 75-mer identifier tags inserted into the bacterial chromosome. This process created roughly 50,000 bacteria that are able to be uniquely identified by the random 7-mer tag with very small (< 2%) risk of duplication in an infection of <50 CFU (See Figure 1 in Martin et al. (25)). The animals were necropsied between 15 and 20 weeks post-infection. Animals were imaged at monthly intervals (or more frequently) to identify timing of granuloma establishment. Pulmonary granulomas were excised during necropsy, and their three-dimensional positions were recorded via matching to PET/CT imaging. Homogenates from excised pulmonary granulomas and infected thoracic lymph nodes were plated, scraped, and sequenced to identify the specific barcode(s) present in each granuloma. Matching the x, y, and z coordinates recorded for each granuloma with its determined barcode content led to a three-dimensional map of the locations of each barcode throughout the pulmonary space. Bacterial burden for each granuloma was determined by counting colonies on the plates.

Three of the four maps are shown in Figure 1 (the fourth was already presented in the original paper (25)). Lung outlines were calculated from terminal scans of each NHP by the process of creating a polygon mesh described above. Small markers represent pulmonary granulomas, while larger markers denote lymph nodes. Each color represents a unique barcode tag. Some samples had more than one barcode tag present, and often these were doublet granulomas (i.e., two granulomas too close in proximity to distinguish at necropsy) and so are marked with a pie chart showing the relative abundance of each barcode tag. The black markers represent pulmonary granulomas for which no barcode tags were found. Filled black markers are granulomas which grew bacteria upon plating but for which barcodes could not be determined, while open markers are granulomas that did not grow bacteria upon plating (sterile); in this study, only CFU+ granulomas were available for barcode determination.

### Model Overview

*MultiGran* is a novel multi-scale, hybrid agent-based model that describes the formation, function, and dissemination of lung granulomas containing Mtb (Figure 2). It uses sampling of nonhomogeneous Poisson processes; rule-based agent placement; parameter randomization; solving systems of non-linear ODEs; and post-process agent groupings to perform *in silico* experiments that track the progress of infection in an individual host. Each granuloma (agent) is placed stochastically within the boundary of the lung environment based on a set of rules. Within each agent, a system of ODEs is linked internally and solved simultaneously to update concentrations of cells, cytokines, and bacterial burdens within each granuloma at every time step. Additionally, within every time step, each granuloma is given the opportunity to disseminate locally and non-locally. *Local dissemination* involves a new granuloma being initialized nearby, while *non-local dissemination* allows initialization anywhere within the lung environment. At the lung scale, the model tracks the development, location, and quantity of granulomas, and determines whether each granuloma is either alone or a member of a larger granuloma cluster. At the granuloma scale, dissemination-event decisions, rules for granuloma formation, and concentrations of all granuloma components are tracked and defined. As is occasionally done when a flexible agent size is needed (35), our agents (granulomas) are placed on a continuous grid. Agents are spherical with dynamically-changing sizes, and granuloma clustering depends on the geometry and position of each of the agents.

**Figure 2:**
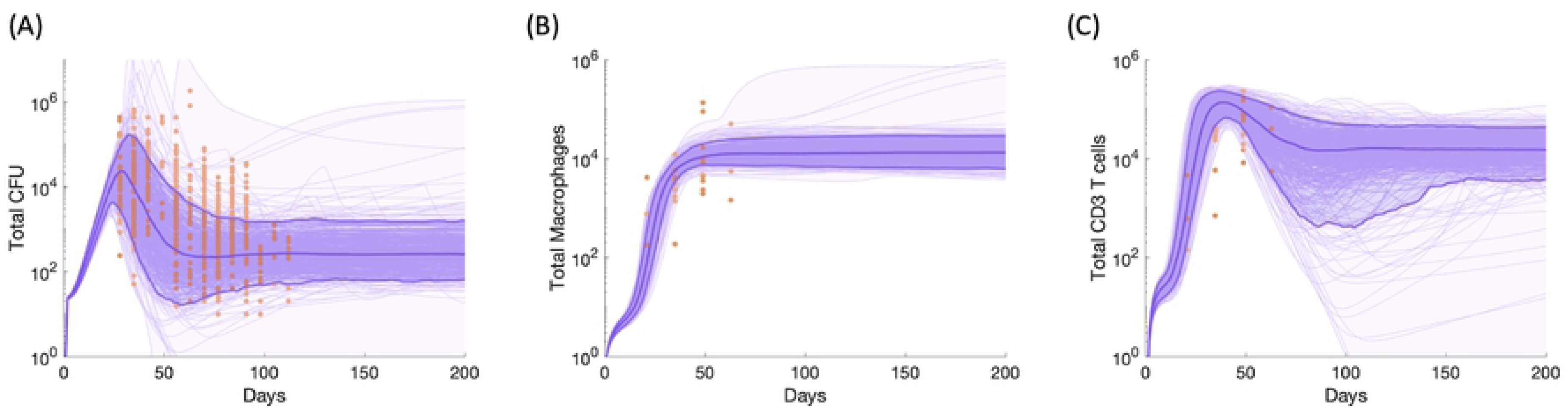
Process of Mtb infection and rules for granuloma dissemination and placement within *MultiGran*. **(A)** A nonhuman primate is inoculated with Mtb, here tracked using different “barcodes” or IDs. These Mtb are taken up by resident macrophages, initiating an innate immune response. This response includes the secretion of various cytokines and chemokines that help prime and/or recruit other immune cells to the site of the infection, resulting in the formation of lung granulomas. Occasionally, as a granuloma develops, it may disseminate--either locally or non-locally. In local dissemination, an Mtb-infected macrophage moves to another nearby location within the same lung lobe. In non-local dissemination, a free extracellular Mtb reaches the airways or is carried to a draining lymph node and then deposited at a site not necessarily near the original location; i.e., in a different lung lobe. Granuloma clusters can form when granulomas develop near each other and may grow into each other, or when one granuloma forms immediately adjacent to the original granuloma via local dissemination (3). Granuloma clusters may contain more than one Mtb ID. **(B)** The rules of granuloma establishment and dissemination within MultiGran. **Case 1 – inoculation.** Inoculation deposits bacteria in a specific lung region at position *(xTrial, yTrail, zTrial)*. The black box designates inoculation region (row 1), wherein the specific within-lung region destined for inoculation is highlighted in green (row 2). The third row demonstrates successful inoculation of a single bacterium – the black box was sampled randomly until the sampled coordinates lie within the green region. Cases 2 and 3 define granuloma placement following dissemination. **Case2 – non-local dissemination**. When non-local dissemination occurs, a bacterium escapes a single granuloma (row 1) and can be placed in any region (shown in black in row 2) that encompasses the entire lung. The green highlighted region is the area in which the bacterial placement will be accepted. Row 3 shows three trial placements: two realizations of accepted bacterial placement (black arrows) and one unaccepted placement (red arrow) at *(xTrial, yTrial, zTrial)*. **Case 3– local dissemination**. Local dissemination is the only form of granuloma placement which does not utilize random placement within a region of lung space. Rather, an infected macrophage from the parent granuloma is placed in a random direction away from the parent granuloma. Row 2 shows several options for granuloma infected macrophage placement. Note that the arrows are of different length to represent our assumption that local dissemination likely follows a normal distribution with respect to parent granuloma location. Here, the green and black arrows show valid directions for the new placement for the infected macrophage, while red arrows show invalid directions. A new granuloma will begin to develop in the chosen (green) valid location (Row 3). Note that in both **(A)** and **(B)** bacteria, granulomas, and infected macrophages are not to scale. Lung image from Servier Medical Art.

Each *in silico* experiment using *MultiGran* is designed to replicate an *in vivo* experiment. To replicate the studies by Martin et al. (25), our simulated NHP is infected with roughly 19 uniquely-identified (barcoded) Mtb that are randomly placed in a localized region of the lungs, similar to the typical inoculation process in the NHP experiments. Each Mtb is assumed to be immediately taken up by a resident lung macrophage, forming a single, unique new granuloma (25). Each granuloma evolves independently. Whenever a granuloma is formed, it is initialized with parameter values that represent several characteristics that ultimately influence its future behavior, as well as the emergent outcomes of the system as a whole.

### Simulation Environment

Code is written in MATLAB, with Bash script for submission to run on computer clusters. ODEs are solved using MATLAB’s ode15s with the NonNegative option for all terms, and we define the start and end time interval to be the size of the agent time step. To avoid complications with the random number generator seed being reset with the initialization of each MATLAB instance, the Bash script executes code that generates a randomized seed list for the simulation to use. The website http://malthus.micro.med.umich.edu/labmovies/MultiGran/ has pseudocode and implementation descriptions, as well as simulation videos.

### Granuloma Establishment

A granuloma is initialized when Mtb is deposited into the lung environment. Based on our previous publications (3, 25), we assume that each Mtb creates one granuloma (3, 36). The granulomas established during inoculation (Figure 2B – Case 1) are referred to as “founder” granulomas and are considered first-generation granulomas; all other granulomas that may emerge throughout the simulation originate from these founders.

Granulomas are agents, so at initialization we assign parameter values to each granuloma and its infecting Mtb, as well as counts and concentrations of all cell types and cytokines. Every granuloma is assigned unique identification markers. These include being given a unique individual granuloma ID *IndivGranID(i)*, which is assigned in chronological order of initialization *i=1,2,…N* (where *N* is the total number of granulomas), as well as the individual granuloma ID of its parent, so the lineage of each of the founder Mtb can be tracked throughout the course of infection. Each granuloma is also given a position on a continuous grid.

### Granuloma Development

The development of each individual granuloma “agent” is captured by a set of ODEs with 16 equations for 16 state variables capturing bacterial, T cell, macrophage and cytokine dynamics (see Appendix 1 for equations and complete term-by-term description of the model). ODE model formulations build on our previous work (37–39) describing cells and levels of cytokines in a whole lung. The equations have been re-calibrated to NHP granuloma data (see section on Experimental dataset) to represent an individual granuloma (see section Model Parameters, Calibration, and Sensitivity Analysis), and have been updated in several ways. First, we increased the role of IL-10, including it as a factor for downregulating macrophage activation and TNF-α production by activated macrophages, as well as allowing infected macrophages to produce IL-10, based on NHP data (40–42). The other set of changes relates to intra- and extra-cellular Mtb to be consistent with recent findings on Mtb growth within macrophages (43–45). Rather than releasing the entire carrying capacity of bacteria at the occurrence of each death of an infected macrophage, the amount of intracellular Mtb within an average infected macrophage is released (with the exception of a bursting infected macrophage, in which case the maximum amount of Mtb is released). Furthermore, only a fraction of intracellular Mtb released during the natural death of an infected macrophage survives to become an extracellular Mtb. The expression for intracellular Mtb replication was also changed along with the addition of an expression for the natural slow death of intracellular Mtb for model stability. We record granuloma sterilization when the count of Mtb drops below 0.5.

### Granuloma Dissemination

While the mechanisms behind dissemination are not yet well-understood (25), we have created rules such that the emergent outcomes are consistent with experiments (Figure 2B). We define a probability function for likelihood of a dissemination event, which we make dependent on the bacterial load (CFU) of the granuloma. We selected CFU because the data presented by Lin et al. (3) indicates that granuloma carrying capacity has a limit (approximately 10^5). Because NHP granulomas rarely exceed this limit (3, 28), there is likely a link between granuloma CFU and dissemination. Because Mtb is by itself non-motile, we consider two routes of dissemination: 1) Mtb conveyance within an infected macrophage and 2) a single Mtb flowing through lung airways or deposited via a draining lymph node (LN). From these, we incorporated two types of dissemination events: local and non-local, the probabilities of each event being independent, and in the unlikely event that multiple dissemination events occur in the same time step, the order of events is randomized.

When a granuloma disseminates locally (Figure 2B – Case 3), an infected macrophage carrying intracellular Mtb is assumed to move from the parent granuloma position to a new position nearby. We assume the distance between the parent granuloma and a new position likely follows a normal distribution with respect to parent location and we calibrated the mean and variance of this location using the data presented in Martin et al. (25). In Martin et al., the authors compute distances of each granuloma and granuloma clusters that they could identify via PET/CT, rather than every individual granuloma regardless of size and cluster affiliation. We also assume that a pre-determined quantity of T cells moves with an infected macrophage. After this dissemination event, the parent and daughter granulomas evolve independently from each other. When a granuloma disseminates non-locally (Figure 2B – Case 2), an extracellular Mtb is simulated as if entering airways (or via a LN) and deposited with equal likelihood anywhere within the lungs, where it is immediately taken up by a macrophage. Figure 2B-Case 2 represents 3 realizations of trial coordinates wherein the trial coordinates represented by the red arrow do not satisfy our criteria, but the two black arrows would be acceptable placements for a bacterium in non-local dissemination.

We created two dissemination event probabilities describing local and non-local dissemination. In both, λ is the maximum probability of dissemination and is scaled by a Michaelis-Menten fraction, using a value of CFU at which the probability is half of the maximum value.

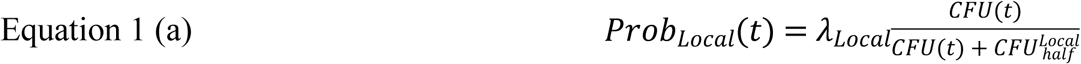

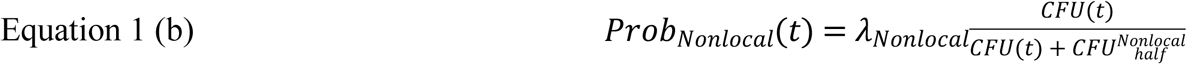

### Granuloma Merging

Experiments demonstrate that a subset of granulomas contain a more than one Mtb barcode (25). Following inoculation or dissemination events, individual granulomas may merge, or are sufficiently close to each other, to form clusters. We identify granuloma clusters and their members when needed for plotting and computing statistics but allow them to evolve independently. Briefly, our algorithm evaluates all intersections of granulomas, and combines groups of granulomas that intersect in 3D space. These grouped granulomas are the granuloma clusters. A granuloma cluster may contain only descendants of a single founder Mtb ID, or may contain descendants of multiple founder Mtb IDs.

### Model Parameters, Calibration, and Sensitivity Analysis

We sought to define the parameter space for *MultiGran* across multiple scales. First, we identified the parameter space of the individual granuloma ODE model that best represents the individual granuloma datasets (CFU and cell counts). To determine an initial, wide range of parameter values to test, we examined experimental values from literature, the previous models (37–39), and values from *GranSim*, our single granuloma model that has been calibrated based largely on NHP data (6–17,19–21). We then used a Latin Hypercube Sampling (LHS) algorithm (46) to sample this multi-dimensional parameter space 500 times. This initial wide range of simulations did not match the NHP data. We narrowed the initial ranges and resampled the space in an iterative process until, out of the 500 simulations, ninety percent of the runs fell within the bounds of our experimental data on CFU, T cell counts, and macrophages within individual NHP granulomas. The parameter ranges for these runs are in Table A1.

Next, we identified the dissemination parameter space of *MultiGran* that matched the NHP whole lung outcome datasets (previously published (20,25,27,31–33) and ongoing studies). We again utilized LHS to sample this space and identify baseline parameter ranges that match the data (Table A3).

Following *MultiGran* calibration, we sampled the calibrated parameter space to create a biorepository of *in silico* lungs that could be used to make predictions and compare to additional NHP data sets. We then used Partial Rank Correlation Coefficient (PRCC), a global sensitivity analysis technique (46), to identify significant correlations between single granuloma ODE model parameter changes and variation in whole lung outputs. We excluded the dissemination parameters from our multi-scale PRCC analysis because they are phenomenological in nature and we are interested in identifying the mechanistic events that occur at the granuloma scale and lead to dissemination, a whole lung outcome.

### Linking Cellular Scale and Tissue Time Scales

We link the cell and cytokine scale events in the ODE model (single granuloma) with the tissue scale ABM (multiple granulomas) to form the multi-scale *MultiGran* model (Figure 2). Linking of timescales is important for proper model design (47). We use an ABM time-step of 1 day. At each ABM time-step, dissemination events can occur. After each ABM time step, the system of ODEs is solved for each granuloma to update the states of all host cells, cytokines and Mtb populations over the next 24 hours. We run the ODEs using adaptive time steps for 1 agent iteration, for each granuloma, before proceeding to the next agent time step, as dissemination events at the agent time step depend on the dynamically-changing state of ODEs. Additionally, the ODE state variable concentrations can be affected by the occurrence of a dissemination event.

## Results

We present a whole lung model, *MultiGran*, that captures the behavior of Mtb infection leading to the development of multiple granulomas via initial infection and then dissemination of bacteria from existing granulomas. We calibrate and validate the model with unique datasets derived from NHPs, the animal model that most closely mimics the features of human infection. We then use the model to identify dissemination rates and to predict mechanisms leading to dissemination.

### Simulated individual granulomas recapitulate in vivo primate granuloma dynamics

We calibrated our single granuloma model, comprised of a system of non-linear ODEs, to data derived from NHP studies. We compared bacterial load (CFU), T cell counts, and macrophage counts over time per granuloma. Our CFU dataset consists of 623 granulomas from 38 NHPs (previously published (20,27,31–33) and ongoing studies). T cell and macrophage counts, as well as additional CFU, were derived from a separate, new dataset of 26 granulomas from 7 Mtb-infected NHPs and baseline data from one uninfected macaque (see Methods). The data from these 7 NHPs capture the timing of the immune system during early events in infection (granulomas from all NHPs were collected between 3-9 weeks post infection) and were imperative for proper calibration of the model.

We identify a range of parameter values (Table A1) that replicate CFU peaks at approximately 35 days and subsequent control of CFU after day 100 post-infection (Figure 3A), macrophage dynamics (Figure 3B), and T-cell dynamics (Figure 3C). These dynamics reflect the initial inability of the innate immune system to control Mtb replication, the eventual control provided by T cells that arrive from the lymph node around day 28, and the stabilization of Mtb counts around day 100. When isolating a suitable parameter range, we identified ranges that matched these overall trends and recapitulated the spread of granuloma outcomes outlined by the NHP datasets. Likely, our spread captures a fuller range of individual granuloma dynamics than a sample from a limited number of NHP can achieve.

**Figure 3:**
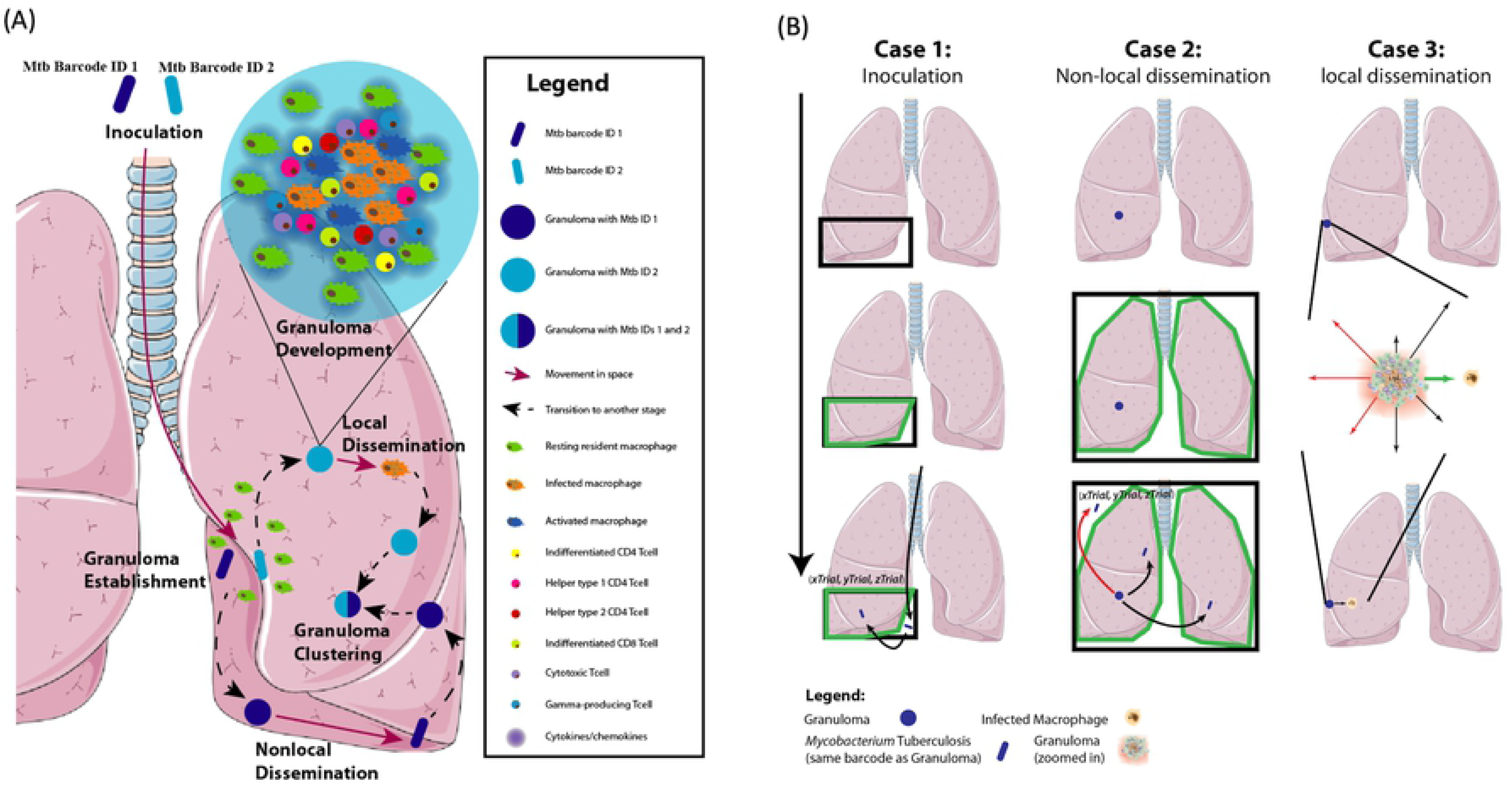
Bacteria, macrophage and T cell dynamics within an individual granuloma. Individual NHP granuloma bacteria **(A)**, macrophages **(B)**, and CD3+ T cells **(C)** shown as orange points across time. Each individual point represents data from a single NHP granuloma. Purple lines indicate simulation outputs from 500 simulations that match NHP data. Light purple shading shows the minimum and maximum of simulation runs, darker purple shading represents the 5^th^ to 95^th^ percentiles of the simulations, and dark purple lines represent the 5^th^, 50^th^, and 95^th^ percentiles of simulations. Parameter ranges are listed in Table A1.

### MultiGran simulates the appearance of granulomas throughout the lung, as seen in vivo

By employing the calibrated single granuloma model (Figure 3) within our *MultiGran* framework, we can now simulate the spread of infection within the lung. We inoculate with 16 to 21 individual bacteria, mimicking the protocol of Martin et al. (25), placing them within an inoculation region within one of the lower lung lobes, as is done in the NHP inoculations via bronchoscope (see Methods). Each initial granuloma in an NHP arises from a single bacterium in an inoculation event (25). Therefore, we initially establish 16-21 granulomas. A sample simulation at the time-point of 250 days post-infection is shown in Figure 4. The blue lung mesh represents the dataset derived from NHPs for (x,y,z) coordinates of a lung. Placed on this mesh are simulation results – individual granulomas (“agents” in the model) and their location, size, and bacterial origin (barcode). Note that, as in the NHP images of Figure 1, infection is primarily within the inoculation region – but that 7 granulomas disseminated non-locally to the opposite lung. In this simulation, one granuloma cluster was found that contained more than one Mtb barcode, as is shown in the pie chart. Movies of disease progression using this 3D visualization are available on the website http://malthus.micro.med.umich.edu/labmovies/MultiGran/.

**Figure 4:**
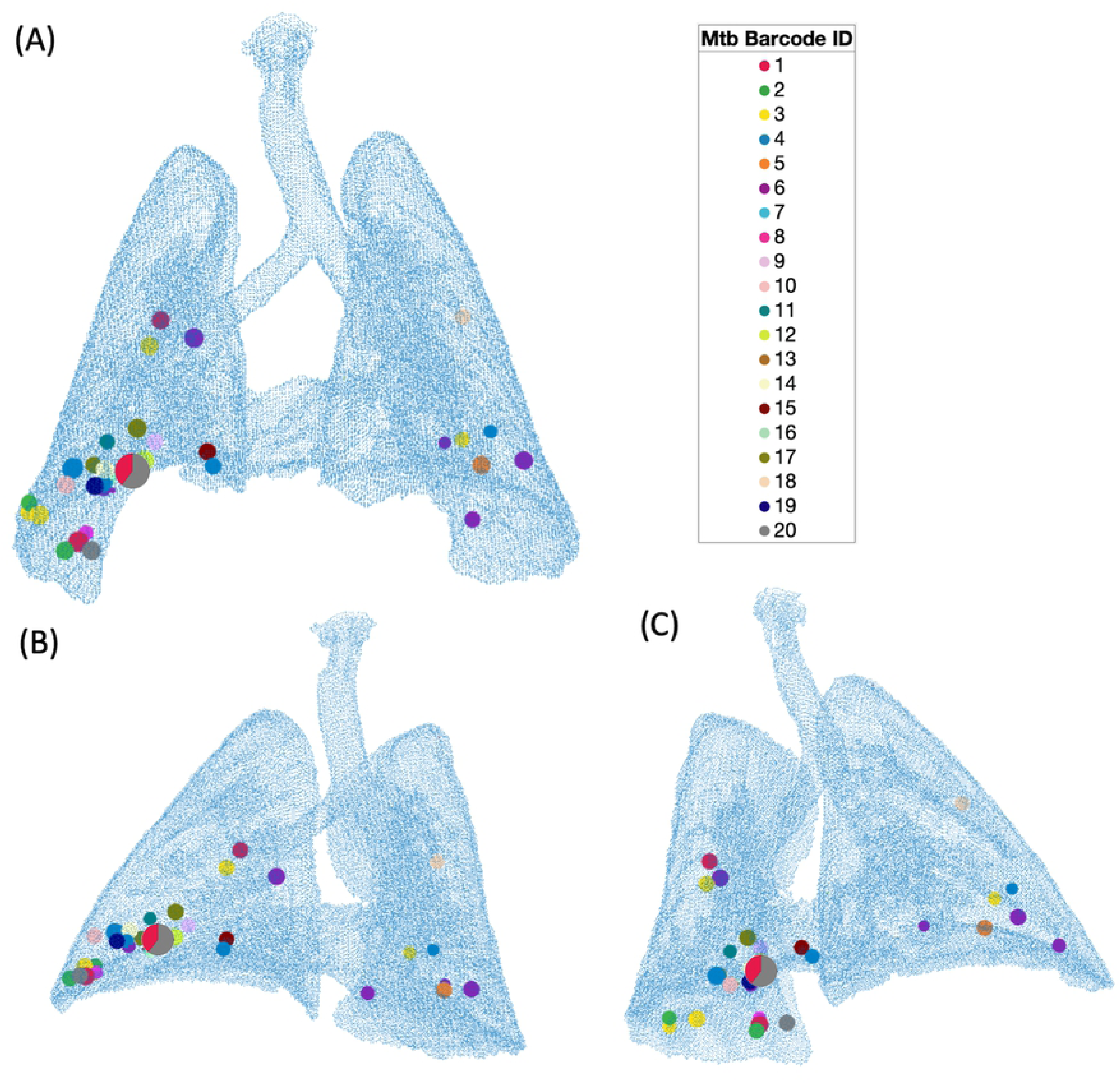
*MultiGran in silico* infection in a non-human primate lung. A single *in silico* simulation at 250 days post infection from three angles (**A**-anterior view, **B&C**-opposite posterior-lateral views), plotted over a data grid taken from PET/CT images of a single NHP. Granulomas are located within the lung in 3D space. Each circle of a single color represents a granuloma or granuloma cluster with a single Mtb barcode ID. The circle shown as a pie chart represents a granuloma cluster with two unique Mtb barcode IDs; each color represents the relative proportion of CFU of each ID compared to the total CFU of the granuloma cluster, while the overall size of the circle is proportional to the size of the cluster. Inoculation was in the lower right lung (bottom left in each image). Granulomas found in the upper right lung and the left lung result from non-local dissemination within the simulation.

### Simulations are consistent with in vivo infection and predict dissemination likelihood rates

*MultiGran* allows both local and non-local dissemination of bacteria to initiate new granulomas, tracks the origin (Mtb ID) of each granuloma, and allows for merging of nearby granulomas to form a cluster. Each granuloma has a unique parameter set chosen from the ranges in Table A1 according to an LHS design. To determine what leads to different dissemination patterns *in vivo*, we use our dataset consisting of four NHPs in that were inoculated with uniquely identifiable Mtb (Figure 1; Martin et al. (25)). Outcome measures from these experiments include: (1) the number of Mtb at time of inoculation (16-21 Mtb), (2) the number of granuloma (or granuloma clusters) at necropsy (17-28 granulomas), (3) the percentage of Mtb barcodes found in multiple granulomas (12.5 - 68.4%), and (4) the percentage of granulomas containing multiple Mtb barcodes (∼10-20%). We calibrated *MultiGran* dissemination dynamics to this dataset by varying the seven dissemination parameters (Table A3). Our whole lung simulations and the NHP dataset are shown in Figure 5. Notice that the simulations capture the full heterogeneity of the *in vivo* results across each NHP. Additionally, the experimental data are from only four NHPs, while our simulations represent a larger, more diverse set of possible outcomes.

**Figure 5:**
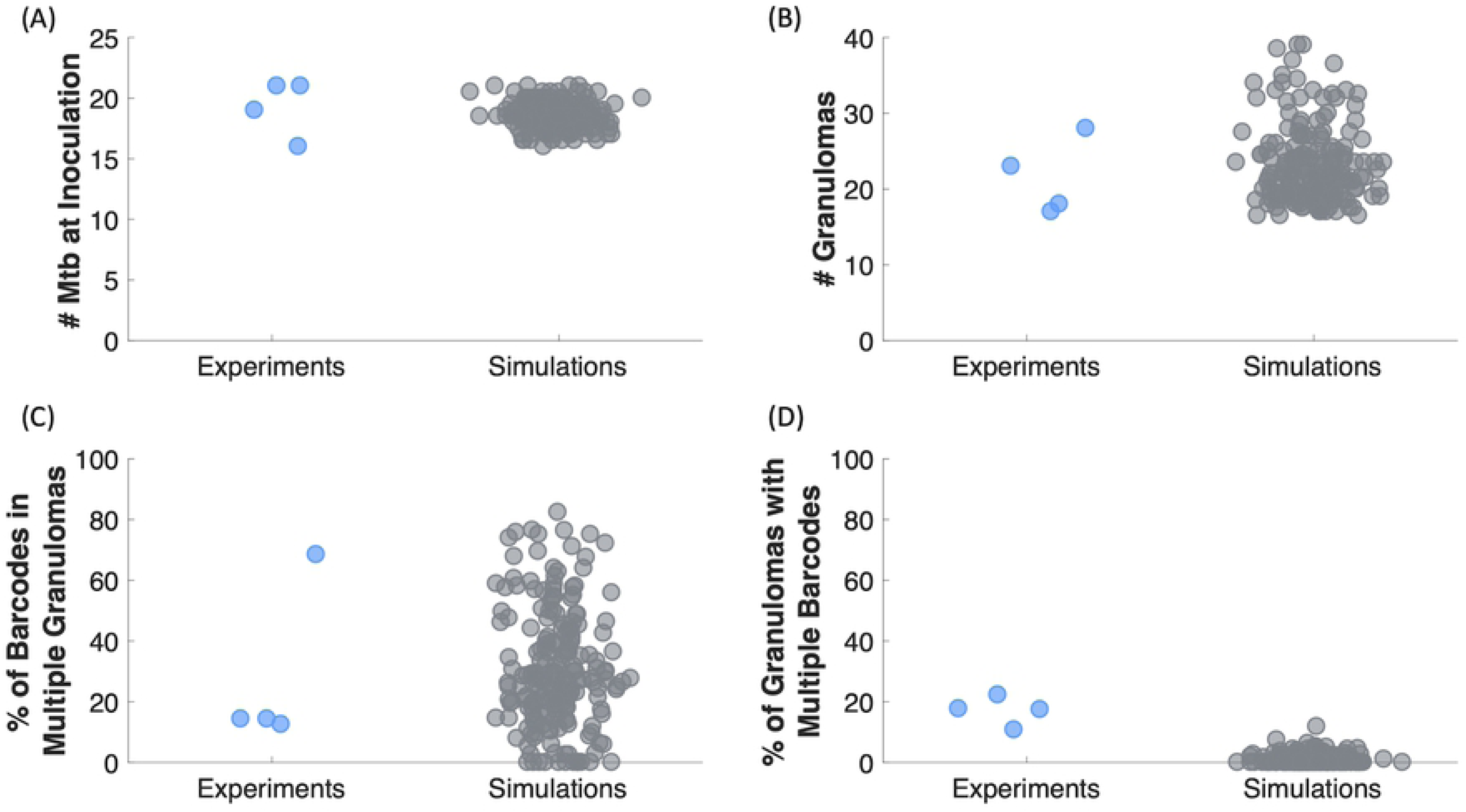
*MultiGran* recapitulates non-human primate dissemination outcomes. Martin et al. (22) infected 4 NHP with 16-21 different Mtb barcodes **(A)**, and after 120 days the NHP immune system formed 16-28 non-sterilized granuloma clusters **(B)**. We replicated these experiments by simulating 200 NHP, which started with 16-21 different Mtb. Of the 16-21 Mtb in NHP, 10%-70% were found in multiple granuloma clusters, meaning at least 10%-70% of Mtb were disseminating. Similar to the NHP data, our simulations have 0%-90% of Mtb barcodes disseminated to multiple granuloma clusters **(C)**. Within the NHP experiments, of the 16-28 non-sterilized granuloma clusters, 10%-25% had multiple Mtb IDs within them, meaning at least 10%-25% of observed granulomas are clusters involving multiple sources of Mtb infection. Our 200 MultiGran simulations demonstrate a similar range of granuloma clusters with multiple Mtb barcodes **(D)**. Simulations are shown in gray whereas NHP experiment outcomes are shown in blue. Each point represents a single NHP or *in silico* simulated granuloma.

To more directly test for non-local dissemination events, we validate our simulations against a second dataset of 38 NHPs (Figure 6). Within this NHP dataset, we identified the lung that contained the most granulomas for each NHP, and termed this lung the more-populated lung. Next, we calculated the percentage of granulomas that resided in the more-populated lung out of the total number of granulomas across both lungs. We found that the 38 NHPs exhibited a range of 52%-100% of granulomas in the lung that was more-populated. Results from the same simulations used to create Figure 5 give a range ∼54%-100%, providing additional support for the model in its ability to capture the range of data offered by NHP experiments.

**Figure 6:**
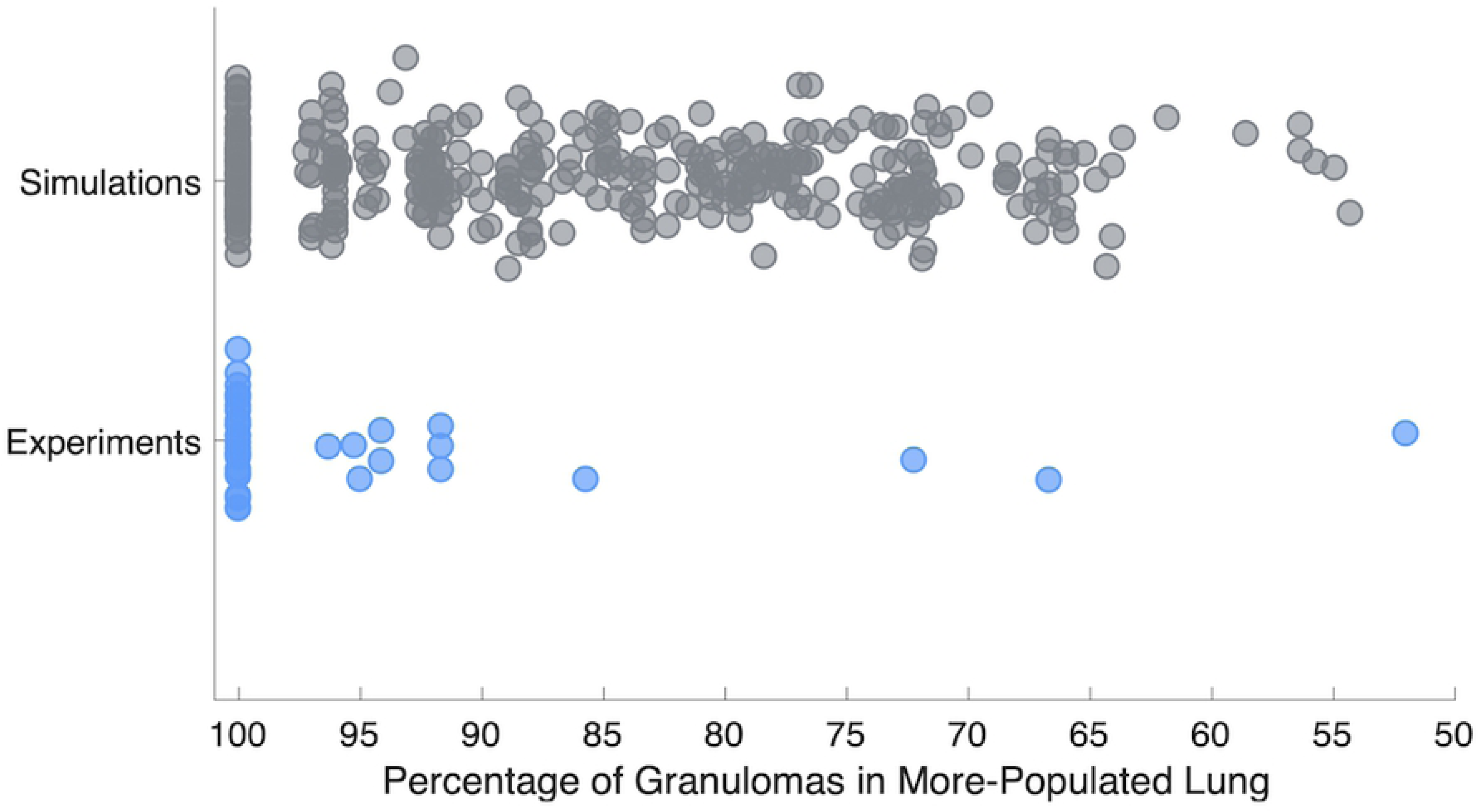
MultiGran recapitulates spread of infection data. At necropsy of 38 NHP experiments, we identified the lung that contained the most granulomas for each NHP. Next, we calculated the percentage of granulomas that resided in the more-populated lung out of the total number of granulomas. We found that 52-100% of granulomas formed resided within the more-populated lung. Blue dots represent each NHP experiment. We ran 200 *in silico* simulations that capture a similar range to the NHP spread of infection from lung to lung, ranging from 54.3% to 100%. Gray dots represent each simulated lung.

When examining *in vivo* data, the total number of dissemination events may be undercounted due to sterilization and granuloma clustering. In contrast, our model is able to count every dissemination event, and thereby provides a predicted frequency of local and non-local dissemination. We found that, on average, the rate of dissemination is about 1/24 dissemination events per granuloma per month for simulations run out to 250 days. Most dissemination occurs earlier in the infection, as noted in Martin, et al. (25). Further, *MultiGran* predicts that local dissemination events occur about twice as frequently as non-local dissemination events.

### MultiGran simulations match individual NHP infections

From our repository of 200 *MultiGran* simulated lungs, we isolated the five simulations that yielded the closest match to the median values of Mtb inoculation (20), the median number of granulomas at necropsy (20.5), the median percentage of Mtb barcodes that were found in multiple granulomas (14.3%), and the median percentage of granulomas that contained multiple Mtb barcodes (17.5%) across the four NHP from Martin et al. (25).

These five simulations represent the best matches to the NHP used in Martin et al. (25). We compare two of these simulations to the CFU/granuloma at necropsy from NHP:179-14 (Figure 7A & 7C). Both lung simulations display satisfactory matches to the NHP CFU data; both simulations cover the spread of the experimental data while lying within the bounds of the dataset. However, while both simulations match the CFU data at 17 weeks, we are able to predict what could have happened beyond the necropsy date by running the simulation for a longer time period. Shown are two distinct possible outcomes with the same parameter set: note they diverge when predicting later dissemination events. Figure 7B shows one simulation predicts bacterial control across all the granulomas within that simulation. Figure 7D shows another outcome. Here, a single granuloma within the lung exhibits uncontrolled bacterial growth leading to dissemination and there is also formation of new granulomas via both local and non-local dissemination (at days 145, 166, and 193). These simulations suggest that NHP:179-14 was either containing the bacteria (i.e., LTBI) (our prediction in Figure 7B) or could have had a subclinical infection that was on the edge of leading to multiple dissemination events (our prediction in Figure 7D). Simulations that match the other NHP are not shown, but show similar trends and predictions.

**Figure 7:**
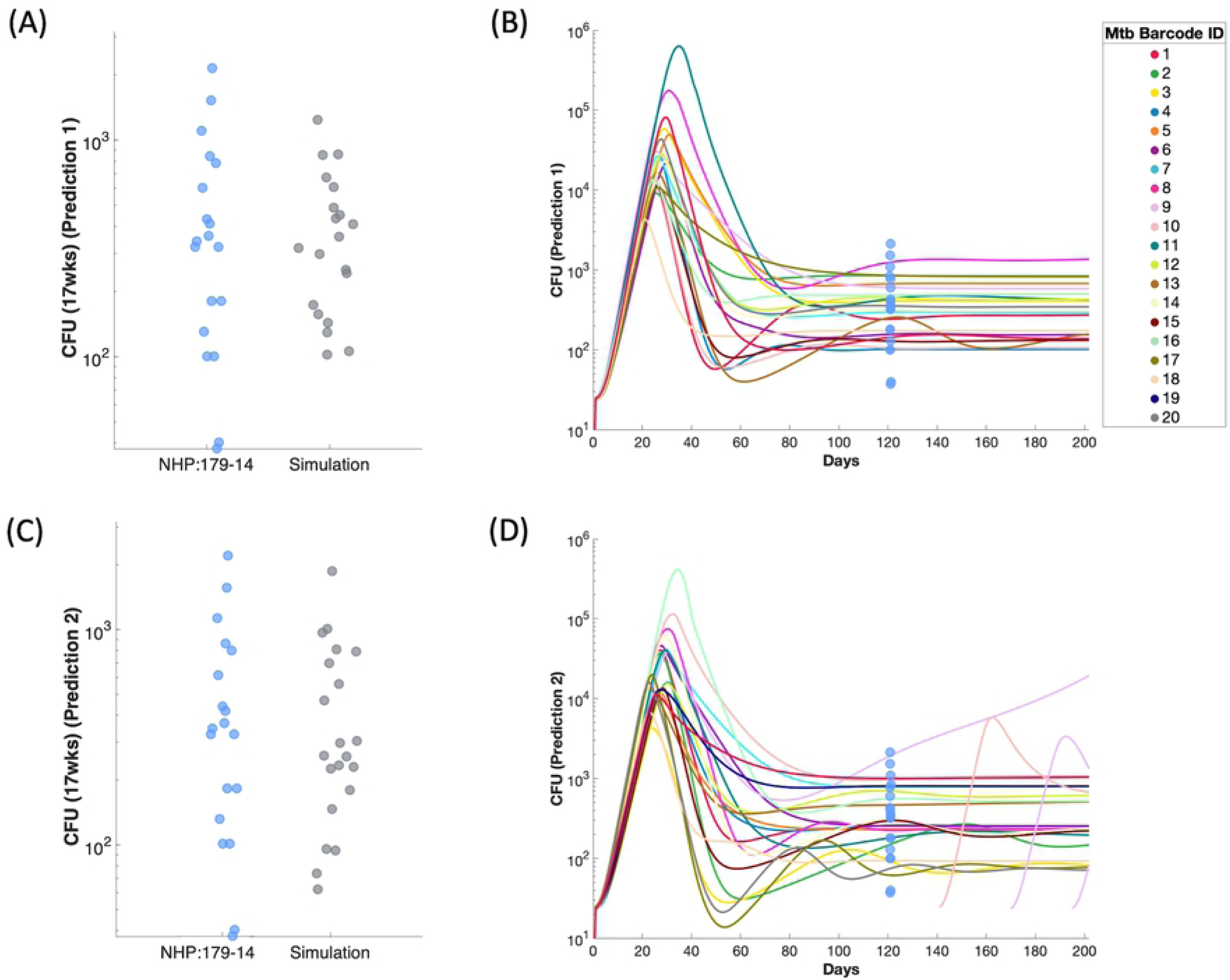
MultiGran matches individual NHP granuloma dynamics and predicts CFU burden across time. We compared the CFU/granuloma at necropsy for NHP:179-14 **(A&C)** to two separate simulations that matched these outcomes. Blue dots represent single granuloma values taken from NHP:179-14; gray dots represent simulation values at comparable timepoints. Simulation predictions diverged after 17 weeks. One simulation predicted stability – i.e., granuloma containment of bacteria **(B)**. The other simulation **(D)** predicted uncontrolled growth of bacteria within one granuloma, leading to dissemination and the formation of other granulomas across time. Each line in **(B&D)** represents one granuloma realization within *MultiGran* across time. Blue dots represent NHP:179 granuloma CFU values. Simulation behavior to the right of the blue dots should be considered a prediction.

### Sensitivity analysis reveals important mechanisms responsible for dissemination

To predict the mechanisms that lead to dissemination events within lungs, we perform global sensitivity analysis on four whole lung outcomes of interest: the number of dissemination events, the total number of granuloma clusters at the end of the simulation, the percentage of granuloma clusters that contain multiple barcodes, and the percentage of granulomas that occupy the initially-inoculated lung at the end of the simulation. We quantify the contributions of each model parameter to the outcomes of interest by calculating partial rank correlation coefficients (PRCC) at the end of the simulation (250 days). Our analysis reveals one parameter as the main driver of these four whole lung outcomes (Table 1). Parameter *CD8MultiFunc* describes the multi-functional nature of CD8+T cells, i.e., the amount of overlap of cytotoxic function and cytokine expression in CD8+ T cells, and is significantly correlated with each of the four outcomes. If *CD8MultiFunc* is increased so that a greater proportion of CD8+ T cells exhibits multi-functionality, then a larger percentage of granulomas will reside within a single lung (less non-local dissemination) and there will be fewer dissemination events and fewer granulomas overall. CD8+ T cells are a key host cell in a functional immune response to Mtb infection, and if the subpopulation that can perform multiple roles within the complex microenvironment of a granuloma increased, it would certainly benefit the host.

**Table 1:** CD8+ T cell functionality plays an important role in dissemination outcomes. PRCCs are shown for parameter *CD8MultiFunc*, the overlap of cytotoxic function and cytokine expression in CD8+ T cells is significantly correlated with each of the four whole lung outcomes at the end of the simulation (200 days). Parameter *CD8MultiFunc* is negatively correlated with the total number of dissemination events across the simulation, the number of granulomas present at the end of the simulation, and the percentage of granulomas that contain multiple barcodes. It is positively correlated with the percentage of granulomas that reside in the more-populated lung.

| Parameter Name | Parameter Description | Number of Dissemination Events | Granulomas at End of Simulation | Granulomas with Multiple Barcodes | Granulomas in Dominant Lung |
| --- | --- | --- | --- | --- | --- |
| CD8MultiFunc | overlap of cytotoxic function and cytokine expression in CD8+ T cells | -0.39 | -0.38 | -0.14 | 0.32 |

If we exclude parameter *CD8MultiFun* from the analysis, we reveal secondary contributions of other parameters to the whole lung outcomes (Table 2). Notably, the role of macrophage-bacteria interactions is found to be important. *k18* represents the base rate of killing of extracellular bacteria by macrophages. If this rate is high, there are fewer dissemination events and fewer granulomas across the simulation. Additionally, *k17* represents the maximum bursting rate of infected macrophages. This parameter is positively correlated with the number of dissemination events and the number of granulomas across a simulation. If bursting occurs at a high rate within a granuloma, our model predicts that a granuloma is more likely to disseminate both locally and non-locally. Taken together, these two parameters identify an important role for macrophage dynamics within the granuloma: if macrophages cannot adequately respond to Mtb, the likelihood of dissemination increases. Altogether, the results of this analysis represent a multi-scale impact: events governing cell function at the cellular scale impact local and non-local dissemination outcomes across the lungs and predict the difference between dissemination and control across the lung environment.

**Table 2:**
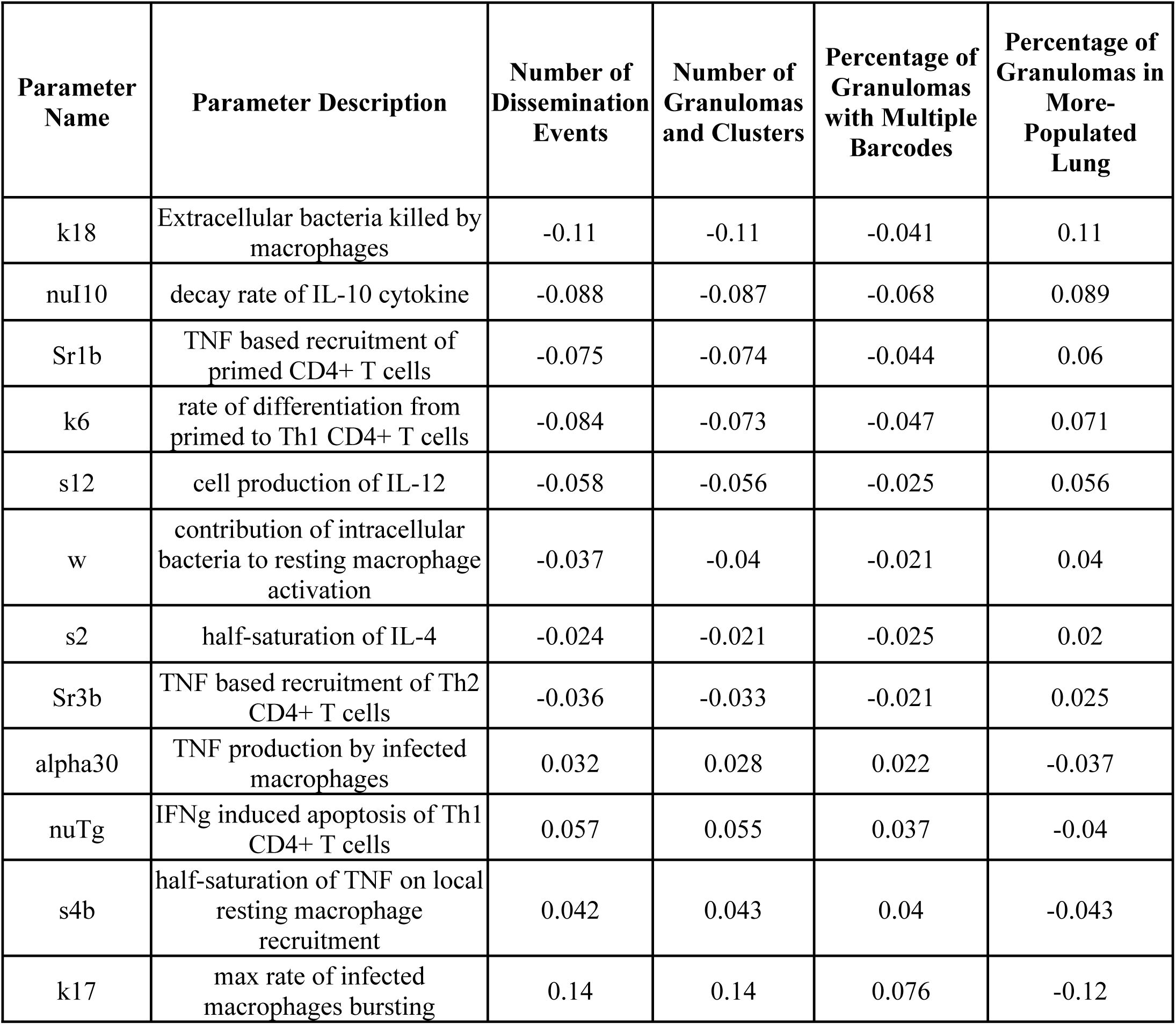
Sensitivity Analysis reveals global drivers of dissemination outcomes. Excluding parameter *CD8MultiFunc*, 12 parameters were identified as having a significant impact on each of 4 *MultiGran* whole lung simulation outcomes at the end of the simulation. All PRCCs shown are significant to p < .05.

## Discussion

Tuberculosis is a complex and heterogeneous disease with a spectrum of outcomes, and the myriad of mechanisms that influence outcomes of initial infection are poorly defined. Our data in NHP models, and bolstered by data in humans, support the notion that each individual granuloma in a host is independent and dynamic, in terms of immunologic composition and function, ability to kill or restrain Mtb bacilli, and risk for dissemination or reactivation (48, 49). However, it can be challenging in NHP models to determine the full range of host mechanisms that play a role in initial containment and prevention of dissemination, both of which are essential to limiting development of active TB. In the pursuit of a better understanding of the collective behavior of lung granulomas in individuals infected with Mtb, we performed a systems biology approach pairing NHP experiments and computational/mathematical modeling. Specifically, we explored events that lead to dissemination and new granuloma formation, and several studies have recently explored this biological phenomenon (25,36,50,51). In particular, the barcoding technique introduced by Martin et al. showed that dissemination varies widely among macaques despite initial infection conditions being similar, and that in individual macaques, each granuloma had a different dissemination risk, from no dissemination by most granulomas, even though these granulomas were CFU+, to multiple dissemination events from a single granuloma. The barcoding analysis provided critical new information about bacterial spread within the lung. However, identifying mechanisms that leading to granuloma dissemination, which is linked to development of active TB (36), is important in designing more effective vaccines and therapeutics against TB. Systems biology approaches can address these mechanisms and more generally contribute to our still limited understanding of Mtb infection dynamics.

In this work, we combine experimental data from NHPs with a novel multi-scale, hybrid agent-based model of granuloma formation, function and dissemination within the lung, called *MultiGran*. We calibrate and validate *MultiGran* against multiple NHP datasets that span cellular, bacterial, granuloma, and whole-lung scales. This calibration and validation allowed us to make predictions about dissemination within Mtb infected lungs. We report that the likelihood of local dissemination is approximately two times greater than non-local dissemination, which supports the in vivo data reported in Martin, et al. (25), and we used sensitivity analysis techniques to identify that dissemination is intertwined with the role of CD8+ T cells in granulomas. Specifically, we predict that the functionality of CD8+ T cells is critically important: if a greater percentage of CD8+ T cells can perform dual functions of cytokine expression (IFNγ, TNF, and IL-10) and cytotoxicity, then the likelihood of dissemination significantly decreases.

The role of CD8+ T cell multi-functionality within the granuloma is controversial (for reviews of CD8+ T cells in TB, see (52, 53), (20)). While the majority of T cells within a granuloma are single cytokine producers (27), multifunctional CD8+ T cells have been demonstrated in the blood of Mtb-infected humans and the proliferation and response rate of these cells differed between active and latent infection (54, 55). Together, these studies and our current work suggest a need for increased focus on this specific cell type to evaluate the potential that CD8+ multifunctional T cells may offer.

The NHP datasets generated within this study are unique and critical to the predictions of *MultiGran*. In addition, these data also present new insights into early events occurring during Mtb infection. In particular, the ability to capture data on Mtb infection during early time points for CFU, T cell counts, and macrophage numbers is instrumental in elaborating timing of early immune response events. These early events in primates have been understudied, and knowledge of the role that timing plays in granuloma establishment, formation, and development is critical to early intervention strategies.

Using *MultiGran*, we were able to match to granuloma population data coming from multiple monkeys (Figure 5 & 6) and granulomas (Figure 3). we were also able to match experimental data from a single NHP (Figure 7). In the era of precision medicine (56), the ability of *MultiGran* to fit to individual data could help predict, in real time, whether the granulomas within that individual are likely to disseminate. This could happen when paired with PET/CT images of individually lung granulomas. However, more realistically, this provides an impetus for identifying biomarkers that are associated with granulomas at risk of dissemination, which could be more widely used to identify persons at risk of developing active TB following infection.

There are a few limitations of our study and model. First, the driving dissemination probability rules are somewhat phenomenological. Our goal in this first study was to rely on as few assumptions as possible; the only granuloma characteristic that is explicitly used in the dissemination rules is the total bacterial burden. As a consequence, the model allows for even a stable, mature granuloma to disseminate (with small probability). We addressed this by allowing T cells to leave the parent granuloma to travel to a daughter granuloma in a local dissemination event, expecting this to sterilize new granulomas. Surprisingly, this was largely ineffective. Instead, it is more likely that the lung parenchyma in infected individuals has increased numbers of Mtb specific T cells and possibly activated macrophages, so that new granulomas form in a completely different immune environment, compared to the initial granulomas that form in an immunologically naïve environment. This notion is supported by our data in NHP models demonstrating that primary ongoing infection protects against reinfection (32). *MultiGran* could be refined to test this in future iterations. Second, we restrict dissemination to be within the boundary of the lungs, but the actual environment within the lungs is very complicated and also could include airways and blood. Third, while we acknowledge thoracic lymph nodes as a source of non-local dissemination, and include adaptive immune cell recruitment in our ODE model, we currently do not explicitly model lymph node compartments. In future work, we plan to address the role of lymph nodes in Mtb infection and dissemination. Finally, while *MultiGran* was developed based on extensive NHP and human data, it does not contain all the various cell types and mechanisms in the complex environment of the granuloma, primarily because the functions and importance of certain cell types and factors remain obscure. As data become available, *MultiGran* can evolve to include additional factors for mechanistic test.

In summary, we utilized a systems biology approach that combined computational modeling and NHP datasets to better understand mechanisms of granuloma dissemination. We present *MultiGran*, the first multi-scale model of granuloma dissemination and formation, that was calibrated and validated to NHP data and we make predictions about the rate of dissemination and the role of specific immune cells in granuloma dissemination. In particular, we discovered roles for multifunctional CD8+ T cells and macrophage dynamics in preventing local and non-local dissemination within the lungs. Altogether, we argue that *MultiGran*, together with NHP experimental approaches, offers great potential to understand and predict dissemination events within Mtb infected lungs.

## Acknowledgments

This research was supported by R01AI123093 awarded to DEK and JLF; R01HL110811 awarded to DEK, JLF, and JLL; and UO1HL131072 awarded to DEK, JLF, and JLL. Simulations also use resources of the National Energy Research Scientific Computing Center, which is supported by the Office of Science of the U.S. Department of Energy under Contract No. ACI-1053575 and the Extreme Science and Engineering Discovery Environment (XSEDE), which is supported by National Science Foundation grant MCB140228. We thank Paul Wolberg for programming assistance. We thank the members of the Flynn lab, especially Nicole Grant, Amy Myers, Mark Rodgers, Jaime Tomko, L. Jim Frye, Brianne Stein, Chelsea Causgrove, Carolyn Bigbee, Pauline Maiello, Alexander White, and Cassaundra Ameel for technical assistance, as well as Dr. Philana Ling Lin and Dr. Charles Scanga for advice and assistance.

## Appendix 1

**(Equations pdf that is attached)**

## Appendix 2

**Table A1:**
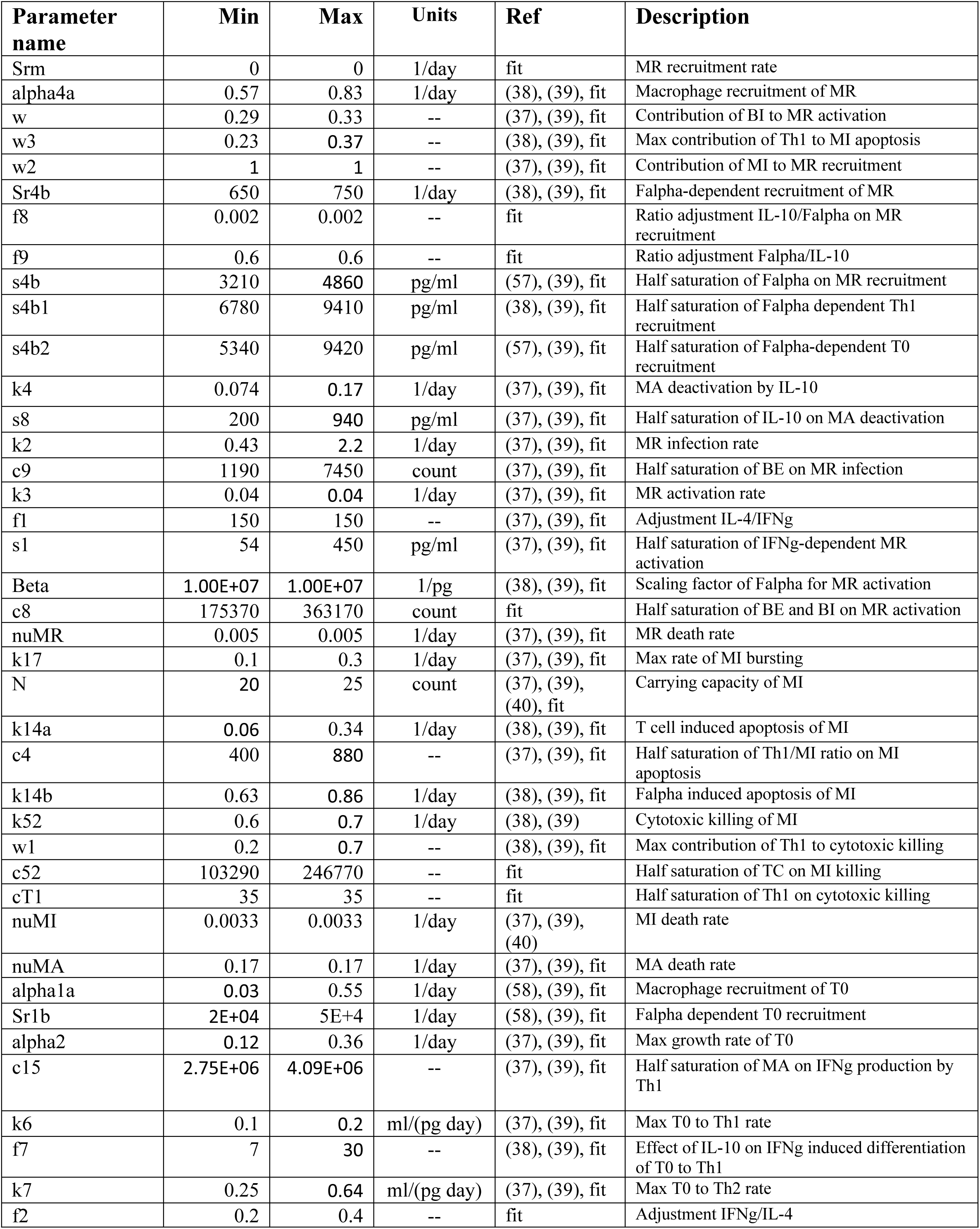
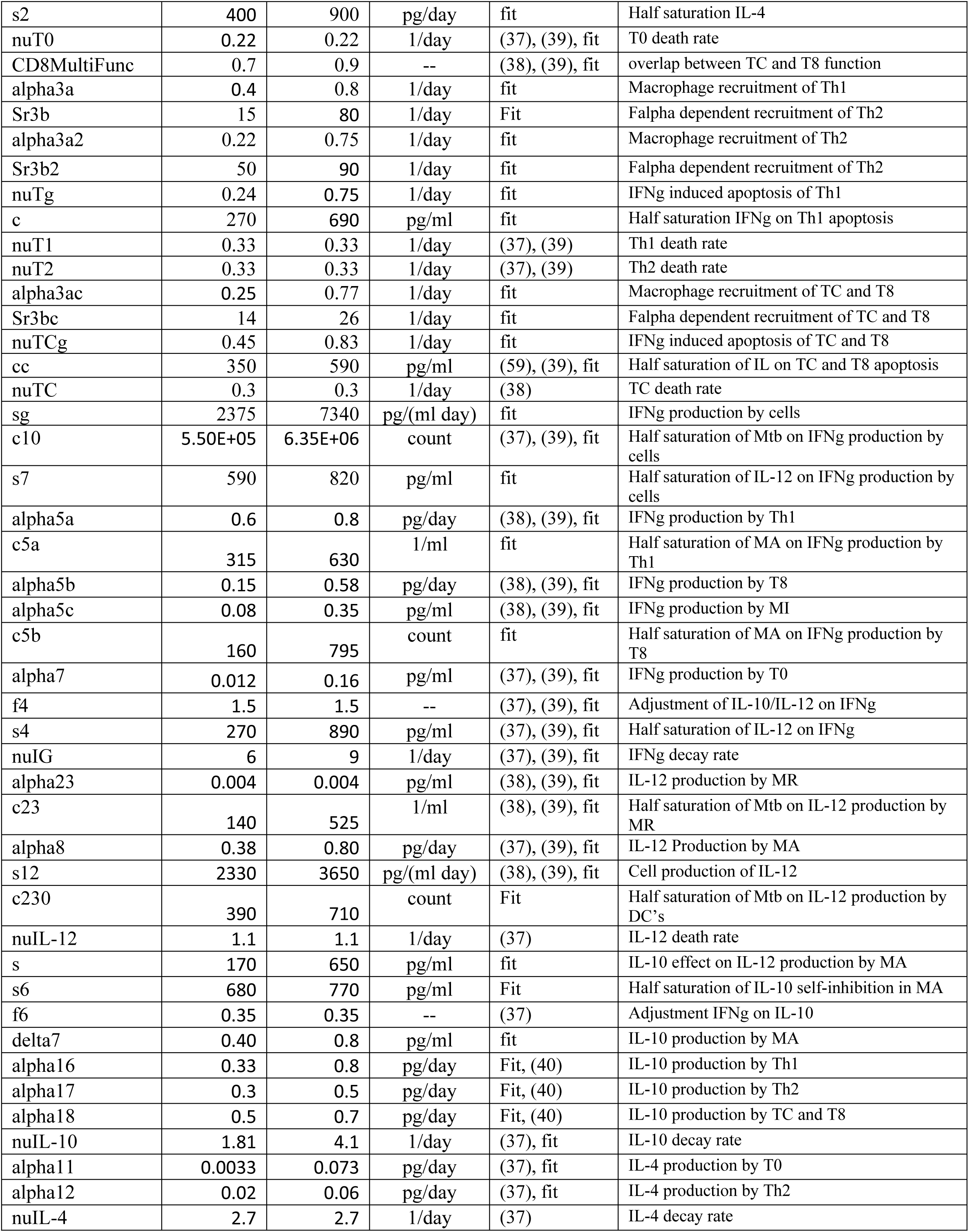
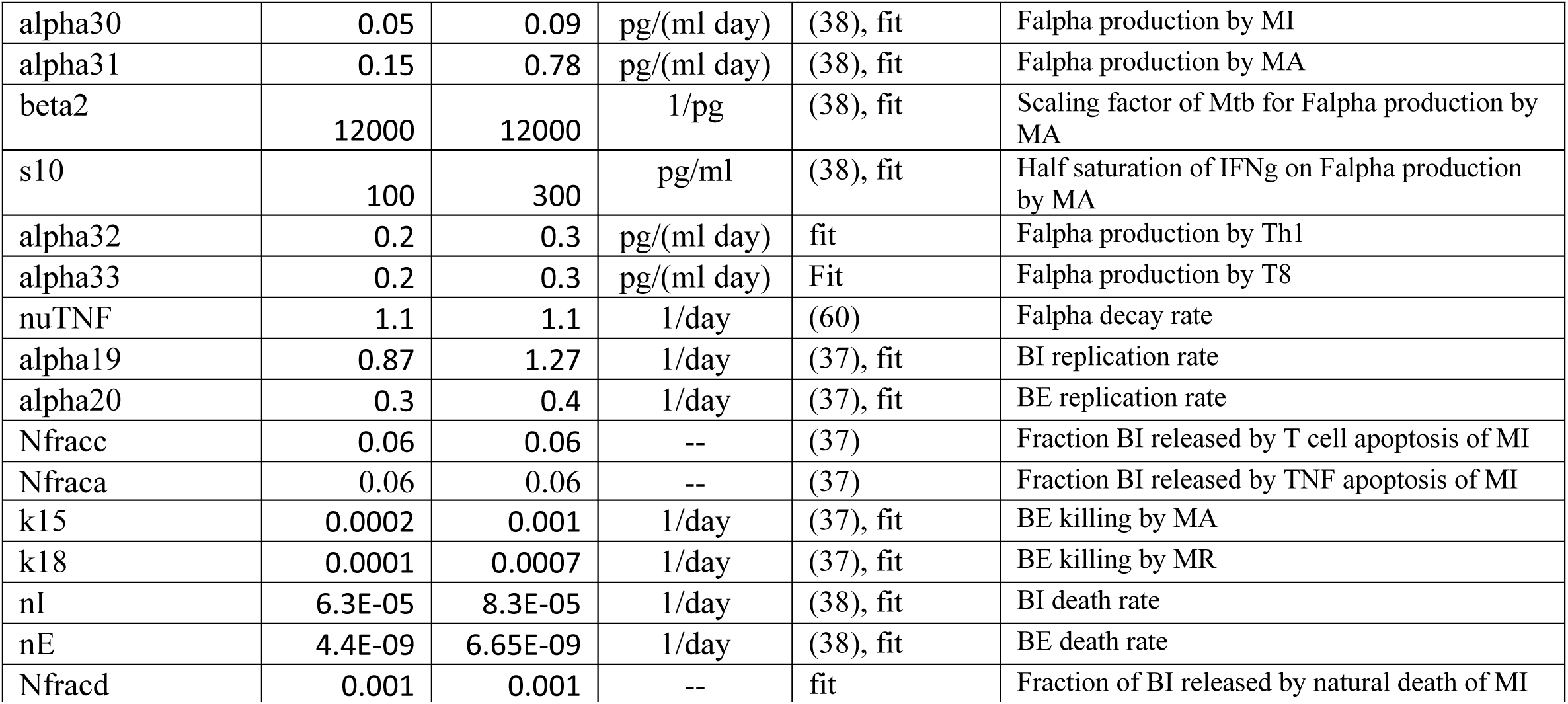
ODE model parameters that govern individual granuloma formation and growth across time. *For each disseminating granuloma, we allow for the option to sample each parameter from a subrange smaller than its parent’s ranges. We do this by using a fraction between 0 and 1 (inclusive) to determine the limits of the range. The fraction represents the percent of values between the parent’s value and either extrema (minimum and maximum) to include in the range. 0 means the range includes only the parent’s value; 1 means that the original range is used.

**Table A2:** Other parameters for size of granulomas and runtime execution.

| Parameter name | Value | Units | Ref | Description |
| --- | --- | --- | --- | --- |
| diamMacs | 20 | microns | (40) | Diameter of Macrophage |
| diamTCells | 5 | microns | (40) | Diameter of T cell |
| dt | 1 | day | ~ | Agent time step |

**Table A3:** Dissemination Parameters. These seven parameters dictate dissemination dynamics in *MultiGran*. Parameters were fit to barcode data or varied using Uncertainty Analysis to find an estimation.

| Parameter name | Min | Max | Units | Reference | Description |
| --- | --- | --- | --- | --- | --- |
| DissemDistMean | $10^0$ | $10^1$ | um | Fit, Based on data (25) | Mean distance of local dissemination |
| Lambda_Local | $10^{-3}$ | $10^{-1}$ | CFU/sec | Fit, Based on data (25) | Max probability of local dissemination |
| CFU_Half_Local | $10^3$ | $10^4$ | CFU | Fit, Based on data (25) | Value for half of max rate of local dissemination |
| Lambda_NonLocal | $10^{-3.5}$ | $10^{-1.5}$ | CFU/sec | Fit, Based on data (25) | Max probability of non-local dissemination |
| CFU_Half_NonLocal | $10^{3.5}$ | $10^{4.5}$ | CFU | Fit, Based on data (25) | Value for half of max rate of non-local dissemination |
| TcellFracDonateMu | 1/100 | 1/10 | -- | estimated | Mean fraction of all of the parent granuloma's T cells that move to the daughter granuloma during a local dissemination event |
| TcellFracDonateSig | $10^{-3}$ | $10^{-2}$ | -- | estimated | Standard deviation from the mean fraction of all of the parent granuloma's Tcells that move to the daughter granuloma during a local dissemination event |

